# The potential for polyphosphate metabolism in Archaea and anaerobic polyphosphate formation in *Methanosarcina mazei*

**DOI:** 10.1101/689885

**Authors:** Fabiana S. Paula, Jason Chin, Anna Schnurer, Bettina Muller, Panagiotis Manesiotis, Nicholas Waters, Katrina A. Macintosh, John P. Quinn, Jasmine Connolly, Florence Abram, John McGrath, Vincent O’Flaherty

## Abstract

Inorganic polyphosphate (polyP) is ubiquitous across all forms of life, but the study of its metabolism has been mainly confined to bacteria and yeasts. Few reports detail the presence and accumulation of polyP in Archaea, and little information is available on its functions and regulation. Here, we report that homologs of bacterial polyP metabolism proteins are present across the major taxa in the Archaea, suggesting that archaeal populations may have a greater contribution to global phosphorus cycling than has previously been recognised. We also demonstrate that polyP accumulation can be induced under strictly anaerobic conditions, in response to changes in phosphate (Pi) availability, i.e. Pi starvation, followed by incubation in Pi replete media (overplus), in cells of the methanogenic archaeon *Methanosarcina mazei*. Pi-starved *M. mazei* cells increased transcript abundance of the PHO-regulated alkaline phosphatase (*phoA*) gene and of the high-affinity phosphate transport (*pstSCAB-phoU*) operon: no increase in polyphosphate kinase 1 (*ppk1*) transcript abundance was observed. Subsequent incubation of Pi-starved *M. mazei* cells under Pi replete conditions, led to a 237% increase in intracellular polyphosphate content and a >5.7-fold increase in *ppk1* gene transcripts. *Ppk1* expression in *M. mazei* thus appears not to be under classical PHO regulon control.

## Introduction

Polyphosphate (polyP) consists of a linear chain of orthophosphate (Pi) residues linked together by phosphoanhydride bonds ranging in length from 3 to greater than 1000 residues^1,2^. It is one of the most widely distributed natural biopolymers, having been detected in many bacteria, fungi, yeasts, plants and animals (reviewed in^3^). Yet despite this ubiquity, polyP metabolism in both the Archaea and Eukaryotes has received scant attention with the majority of studies focussing on bacterial polyP metabolism. These have proposed a range of biological roles for polyP including serving as Pi and energy reserves during periods of nutrient limitation or environmental change, and enabling resistance to environmental stresses such as heavy metals^4–6^, ultraviolet radiation^7^, salinity^8^, pH^9,10^, and exposure to heat, sulphide and other oxidants^11^. Additionally, polyP has been associated with a multitude of other physiological functions including the regulation of gene expression, modulation of enzyme activity, biofilm formation, signalling, virulence, and cation sequestration and storage^12,13^.

Bacterial polyP synthesis is primarily catalysed by the enzymes polyphosphate kinase 1 (PPK1) and 2 (PPK2), which transfer the terminal Pi residue of a nucleoside triphosphate (ATP for PPK1, and GTP or ATP for PPK2) onto a growing chain of polyP. Biopolymer degradation is catalysed predominantly by exopolyphosphatase (PPX), which hydrolyses terminal Pi residues^2,14^.

Several ultrastructural investigations have observed the presence of polyP granules in archaeal isolates e.g. *Sulfolobus* sp., *Archaeaglobus fulgidus, Metallosphaera sedula, Methanospirillum hungatei* and in members of the *Methanosarcinaceae*^13,15–17^. In some archaea, polyP accumulation has been shown to be an important resistance mechanism against metal toxicity^5,6^, and exposure to oxidative stress^18,19^. Orell *et al.*^20^ also noted the presence of homologs of bacterial *ppx* in 9 members of the Crenarchaeota and in 1 Euryarchaeota, and *ppk* homologs were identified in 9 Euryarchaeota, whilst Zhang *et al.*^21^ reported the presence of *ppk1* and / or *ppk2* homologs in *Methanosarcina acetivorans, Methanosarcina mazei* and *Haloferax volcanii*: PPX activity has also been characterised in cells of *S. solfataricus* where polyP accumulation plays a role in resistance to copper stress. Despite this, a focussed study of archaeal polyP metabolism has yet to be carried out, and so our understanding of the enzymology, gene regulation and environmental triggers for polyP metabolism in the Archaea lags behind that of the Bacteria.

Wang *et al*.^22^ have previously analysed 427 Archaeal reference proteomes for the pathways of polyP metabolism and identified PPK1, PPK2 and PPX within the genera *Methanolobus, Methanoregula*, and *Methanosarcina*. In this paper we interrogate the RefSeq database which, at the time of examination (November 2018), consisted of sequences from 1,126 Archaea to further investigate the distribution of polyP metabolism proteins across the Archaeal domain. Moreover, we investigated how polyP accumulation in cells of the methanogenic archaeon *Methanosarcina mazei* could be induced by changes in Pi availability. Using a combination of chemical phosphorus (P) analysis, fluorescence microscopy and transcriptomics, we assessed physiological changes in *M. mazei* under conditions of “polyP overplus” i.e. Pi starvation, followed by incubation in Pi replete (overplus) media: Overplus is the classical experimental model for the investigation of polyP accumulation in bacteria and yeasts^23^. This study is the first to report the phenomenon of polyP overplus in a methanogenic archaeon, and the occurrence of overplus under strictly anaerobic conditions. With this approach, we not only assessed the cellular regulation of polyP turnover, but also explored the responses of *M. mazei* to Pi oscillations. Methanogens have a key role in carbon cycling in anoxic environments and thus studying their adaptive responses to Pi availability may reveal novel aspects of their unique ecological niche and factors controlling their activity. A further understanding of the distribution and regulation of polyP turnover in archaea will not only provide insights into the ability of these microorganisms to tolerate extreme conditions but will also extend our understanding of P cycling in low redox natural and engineered environments.

## Methods

### Database searches and phylogenetic analysis

PPK1, PPK2 and PPX query sequences were obtained from the National Center for Biotechnology Information’s (NCBI) Protein database (**Table S1**). The RefSeq database was queried using these proteins and BLASTP^24^, restricted to the domain Archaea, and with the hit limit set to 20,000. Hits for each reference sequence with an e-value of 1e^-10^ or lower were collated and duplicates removed using the dedupe2 tool in BBTools (Joint Genome Institute, Lawrence Berkeley National Laboratory, USA: https://jgi.doe.gov/data-and-tools/bbtools/). The lengths of the sequences were compared and any sequence with a length less than half of the average was removed in order to eliminate partial sequences. Unique sequences were combined with one of the bacterial reference sequences (Proteobacterial PPK1 [WP_000529576.1] for PPK1, *Pseudomonas aeruginosa* PPK2 [WP_003112634.1] for PPK2 and *Escherichia coli* PPX [EDU80702.1] for PPX) and aligned by MUSCLE^25^ using the default parameters. The alignment was trimmed using TrimAl^26^ under a range of settings, including *automated1, gappyout, noallgaps, nogaps, strict*, and *strictplus*, as well as the settings used by Luo *et al*.^27^ (*-resoverlap 0.55-seqoverlap 60*). Each of the resulting alignments was processed by IQ-TREE^28^ using ModelFinder^29^ with 1,000 bootstraps^30^. The tree with the highest minimum bootstrap value was used to produce a cladogram with the Interactive Tree of Life^31^. Putative identification of the taxonomic order which the organism possessing each homolog belonged to was performed by comparing the organism identifier of the sequence FASTA header to the NCBI taxonomic database.

### Growth conditions

The archaeal strain *Methanosarcina mazei* S-6 DSMZ 2053 (DSMZ, Braunschweig, Germany) was used throughout this study. *M. mazei* was grown in bicarbonate-buffered basal medium (BM)^32^ with the modifications described before^33^. Briefly, the initial mixture contained the following components, in g L^-1^: KH_2_PO_4_, 0.41; Na_2_HPO_4_, 0.43; Na_2_SeO_3_.5H_2_O, 0.3; and Na_2_WO_4_.2H_2_O, 0.3. The medium was complemented with yeast extract (0.2 g L^-1^), boiled down to a final volume of 900 mL (approx. 20 min) and cooled while being flushed with N_2_. After cooling, the medium was dispensed into bottles (360 mL per bottle), while being flushed with N_2_. The bottles were sealed, and the headspace gas was changed by evacuating to −1 atm pressure and pressurising to 0.2 atm with N_2_/CO_2_ (80/20 v/v) three successive times in order to obtain an oxygen-free environment. After autoclaving, the following sterile solutions were added: mixture C1 (50 mL L^-1^); and mixture C2 (50 mL L^-1^), yielding a final pH of 7.0–7.2. The mixtures C1 and C2 were prepared as described before^33^: C1 contained trace metal, vitamin, and salt solutions, and mixture C2 contained NaHCO_3_ (80 g L^-1^), Na_2_S.9H_2_0 (240.2 g L^-1^), and cysteine-HCl (10 g L^-1^). The medium was supplemented with sodium acetate (2.5 g L^-1^) and methanol (5 mL L^-1^). The bottles were inoculated with pre-grown cultures (1:10), and then incubated in the dark at 37 °C without shaking.

### Pi-starvation / overplus experiment

*M. mazei* was grown for seven days as described above. Thereafter, 18 mL of culture was transferred anaerobically into round bottom 43 mL sterile glass vials (Glasgeraetebau Ochs, Bovenden, Germany), with N_2_/CO_2_ atmosphere. The Pi-starved/overplus treatment was carried out as follows: to establish the Pi-starvation phase, the following steps were carried out twice: 1) the glass vials were depressurised and centrifuged at 3000 x g for 40 min, at 15°C; 2) the vials were re-pressurised to 0.2 atm, and the medium was carefully removed through a double-ended needle, coupled to a flask with negative pressure; 3) the cells received 18 mL of fresh medium, prepared as described above (including carbon source), but lacking the phosphate components (KH_2_PO_4_ and Na_2_HPO_4_). The absence of the phosphate buffer did not affect the pH of the medium, which was kept within the original range by the bicarbonate buffer in the mixture C2. Pi-replete control vials were processed with the same steps described above but received full medium containing Pi during step three. Pi-starved and Pi-replete control cultures were incubated in the dark at 37 °C without shaking for four days (days 0-4). On day 4, the starved cultures received 6 mM of phosphate (equimolar KH_2_PO_4_ and Na_2_HPO_4_), initiating the Pi-overplus phase. Both treatment and control cells were incubated for additional two days (days 4-6). Extracellular Pi concentration, protein content and CH_4_ production were monitored daily from triplicate vials from both treatment and control. Destructive sampling was performed in triplicate on days 4 (T4, Pi-starvation sampling time) and 6 (T6, Pi-overplus sampling time) to assess gene expression in both treatment and control cells.

### Analytical methods

To monitor methane production, 1 mL of headspace gas was analysed by GC using a Clarus 500 Gas Chromatograph equipped with a TurboMatrix 110 Headspace sampler (Perkin Elmer Ariel, Waltham, MA, USA). The gas volume was corrected by measuring the expanded volume of 1 mL sample in the 2 mL syringe.

To measure Pi concentration in the medium and cell protein content, 0.5 mL of culture was centrifuged at 4000 x g for 3 min. Pi concentration in the supernatant was assessed by colorimetric method, using the BIOMOL Green kit (Enzo Life Sciences, Farmingdale, NY, USA) and a PowerWave HT spectrophotometer (Biotek, Winooski, VT, USA), according to the manufacturer’s instructions. Protein concentration, used as a proxy for biomass, was estimated with a DC™ Protein Assay Kit II (Bio Rad Laboratories, Hercules, CA, USA). Cell pellets were lysed by adding 200 µl of the kit’s solution ‘A’, followed by vortex homogenisation for 8 min, and heating at 60 °C for 1 min. The protein quantification in the lysate was performed according to the manufacturer’s instructions.

To extract polyphosphate from overplus and control samples, cultures were grown and phosphate stressed as described above. After 4 or 6 days 18 mL volumes were transferred to 50 mL conical tubes and flash frozen with liquid nitrogen. Samples were defrosted on ice and centrifuged at 10,000 x g and 4°C for 1 minute and the supernatant discarded. The pellets were resuspended in SDS lysis buffer (0.5 mL 5% w/v SDS, 50 mM EDTA, 125 mM pH 7.4 Tris buffer) and transferred to a Lysing Matrix E tube (MP Biomedicals, Santa Ana CA, USA), with 500 µl 25:24:1 phenol/chloroform/isoamyl alcohol. The cells were lysed using a FastPrep 120 (ThermoFisher Scientific, Paisley, UK) for two bursts of 30 s at 4.5 m s^-1^. The tubes were centrifuged at 18,000 x g and 4 °C for 15 min and the aqueous phase transferred to a 1.5 mL Eppendorf tube (Eppendorf UK Limited, Stevenage, UK). Two washes with 500 µl 24:1 chloroform/isoamyl alcohol were performed, with centrifugation and retention of the aqueous phase as above. Ice cold ethanol (2.5 vols) and 175 µl of 3 M sodium acetate solution were added and the tubes incubated overnight at −20 °C. The tubes were centrifuged as above and supernatants discarded. The pellets, containing precipitated polyanionic molecules including polyphosphate and DNA, were washed with 1mL 100% and 70% ice cold ethanol and centrifuged as above, air-dried for 15 minutes and then resuspended in 50 µl nuclease-free water. Polyphosphate was detected using a modified version of the protocol described by Werner *et al.*^34^, where polyphosphate is hydrolysed by recombinant *Saccharomyces cerevisiae* PPX and the released Pi quantified. Assays were carried out in 50 µl volumes containing 10 µl of extract, 5 µl nuclease-free water, 10 µl 2 mg ml^-1^ ScPPX and 25 µl 100 mM pH 7.4 Tris buffer with 10 mM MgCl_2_. Assays were run in triplicate overnight at 37°C and 200 RPM shaking, alongside polyP-free and protein-free controls Phosphate release was detected using the molybdate method described by Werner *et al.*^34^ with 20 µl of sample in a transparent 96-well plate and a Fluostar Omega plate reader (BMG Labtech, Buckinghamshire, UK).

### DAPI staining and fluorescence microscopy

Culture samples (1 mL) were withdrawn with a syringe and transferred into 2 mL anaerobic vials, containing N_2_/CO_2_ atmosphere. The cells were fixed with formaldehyde 4 % (in phosphate saline buffer, PBS; pH 7.2), for 15 min at room temperature. Subsequently, the samples were centrifuged at 4000 x g for 3 min and washed three times in PBS. The fixed cells were stained with 50 µg mL^-1^ of 4’,6-diamidino-2-phenylindole (DAPI; Sigma-Aldrich, St. Louis, MO, USA) and incubated for 1 h, at 4 °C. The stained cells were washed twice in saline solution (NaCl, 0.85 %) and resuspended in distilled water. Cell suspensions (5 µl) were spread on glass slides, air dried and analysed under 100 x/1.3 oil objectives by an Axio Imager M2 epifluorescence microscopy, coupled to AxioCam IC camera (Carl Zeiss Vision Inc., San Diego, USA). Simultaneous detection of three fluorophores Blue/Green/Red was carried out with the filter set 62 HE BFP + GFP + HcRed shift free (E) (Carl Zeiss Vision Inc.). Three band pass filters were applied for excitation (350-390, 460-488, 567-602) and emission (402-448, 500-557, 615-4095). Images were analysed with the software ZEN 2.3 (Carl Zeiss Microscopy GmbH, 2011).

### Gene expression analysis

Transcriptomic analysis was performed using triplicate vials from the Pi-starved/overplus treatment and the Pi-replete control, collected at days 4 (Pi starved) and 6 (Pi-replete). Total RNA was purified using ZR Soil/Fecal RNA Kit (Zymo Research, Irvine, CA, USA), with the modifications described in the ref.^35^. Samples were DNase treated as recommended in the Zymo Research manual. Ribosomal RNA was depleted using Ribo-Zero rRNA Removal Kit (Illumina, San Diego, CA, USA), according to the manufacturer’s recommendations. Quantity and quality of total RNA and depleted RNA samples were assessed using the RNA 6000 Nano Lab Chip Kit on a Bioanalyzer 2100 (Agilent Technologies, Santa Clara, CA, USA). Library preparation and sequencing were performed by SciLife (Stockholm, Sweden). Briefly, for each sample, 80 ng of rRNA-depleted RNA was used to prepare paired-end libraries, constructed using the Illumina TruSeq Stranded mRNA Library Prep Kit (Illumina), on an Agilent Bravo Automated Liquid Handling Platform (Agilent Technologies, Santa Clara, CA, USA). The normalised libraries were sequenced on an Illumina MiSeq platform (Illumina). We obtained a total of 9.6 million sequences from the Pi starved transcriptome (T4), 9.0 million from the overplus cultures (T6) and 9.9 and 10.2 million sequences in the Pi replete control transcriptome at T4 and T6, respectively, with mean paired-end sequence length of 152 bp. Transcriptomic data analyses were performed using Galaxy^36^ and R 3.4.4^37^ platforms. Reads passing quality control (according to Galaxy’s FASTQC standard protocol) were mapped to the genome of *M. mazei* S-6 (NZ_CP009512.1) using Bowtie2^38^. Approximately 98% of the transcripts were mapped to coding sequences of the *M. mazei* genome. To quantify gene expression, the number of reads unambiguously mapped per gene was counted with featureCounts^39^. DESeq2^40^ was used to normalise counts across all samples and to compare relative gene expression between treatments or sampling times, with two factor levels. The false discovery rate (FDR) control method was applied to correct *p* values, due to the multiple tests performed^41^. Genes with log2 fold change >2 and FDR< 0.05^42^ were defined as differentially expressed. To evaluate enriched functions, gene ontology (GO) terms for genes up- and down-regulated under overplus conditions, in comparison to Pi starved cells, were retrieved from Uniprot and summarised using REVIGO^43^ (details in supplementary material). Sequencing data has been deposited in the Sequence Read Archive of the National Center for Biotechnology (accession numbers SAMN08132863 - SAMN08132874).

## Results

### PPK1 PPK2 and PPX homologs in Archaea

In total 442 homologs of PPK1, PPK2 and PPX were found in Archaeal sequences of the RefSeq database. Alignments trimmed with the *nogaps* (PPK1) and *strictplus* (PPK2 and PPX) methods produced cladograms with the highest bootstrap values.

Searches for archaeal homologs to the PPK1 reference proteins returned 242 unique sequences (**Figures 1A and S1**). Of these, 241 sequences were identified to at least the order level in the NCBI taxonomic database distributed across eight different orders, all belonging to the phylum Euryarchaeota. The most represented orders were the Haloferacales, Methanosarcinales and Halobacteriales with 44.6%, 31.5% and 12.1% of the homologs respectively, while the remaining identified sequences were from the orders Natrialbales (12.0%), Methanomicrobiales (7.0%), Methanomassiliicoccales (1.7%) or Thermoplasmatales (0.4%). PPK1 homologs from methanogenic archaea formed three major clades, which were distantly related to sequences from the halophilic archaea (orders Halobacteriales, Haloferacales and Natrialbales).

**Figure 1.**
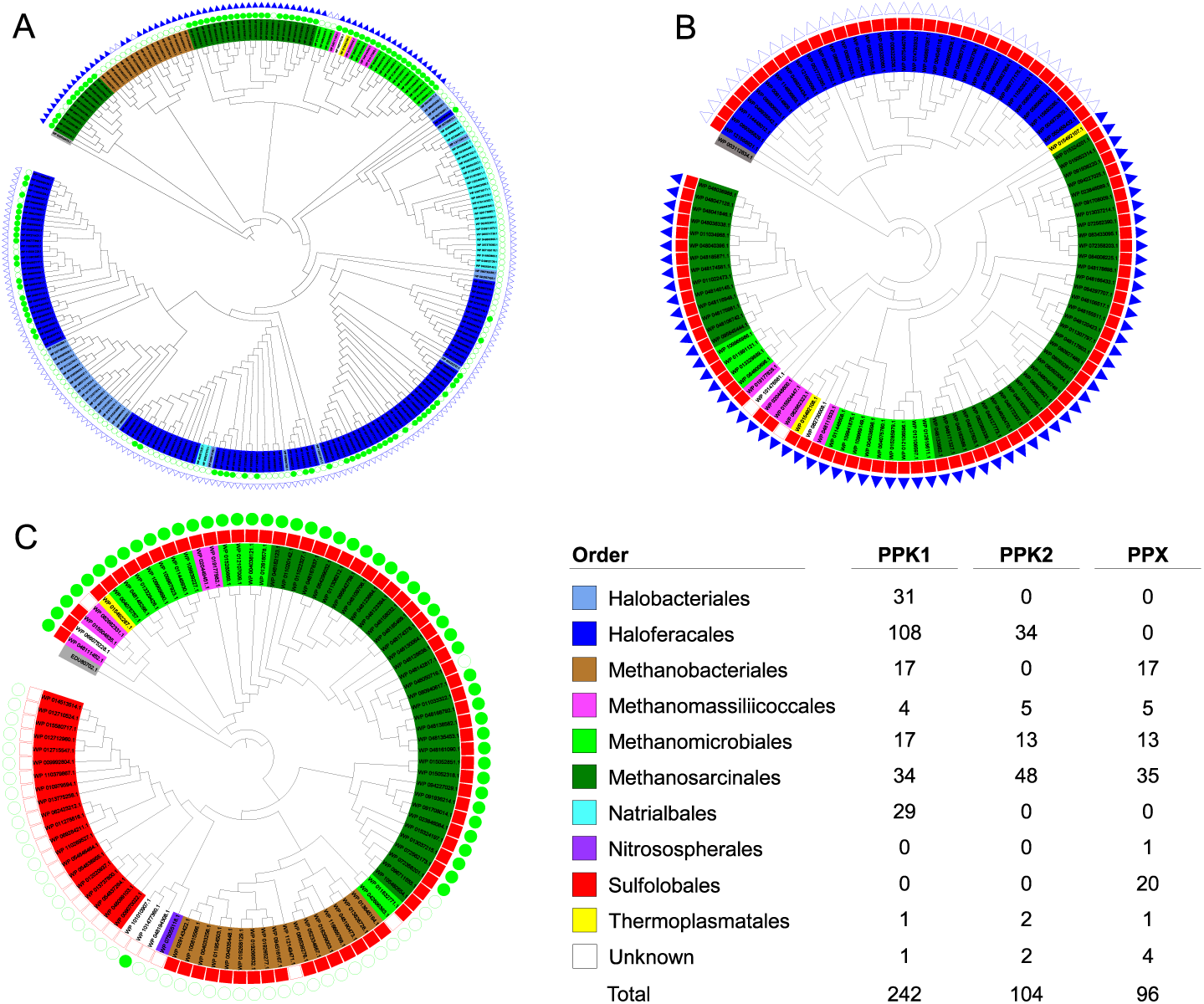
Cladograms of archaeal homologs of PPK1 (A), PPK2 (B) and PPX (C) proteins. Branch tips represent protein homologs and are coloured according to the order which the sequence belongs to according to the NCBI taxonomic database. Values in the table refer to the number protein homologs retrieved from the database. Symbols outside the cladograms indicate whether (filled symbol) or not (empty symbol) a sequence from an organism with an identical name was found in the PPK1 (red squares), PPK2 (green circles) or PPX (blue triangles) figures for comparison.

104 archaeal PPK2 homologs were found (**Figures 1B and S1**). All but two sequences were identified to at least the order level and were distributed across five different orders. The sequences largely originated from the methanogens, with the Methanosarcinales, Methanomicrobiales and Methanomassiliicoccales representing 46.2%, 12.5% and 4.8% of the homologs, respectively. As for PPK1, halophilic archaeal PPK2 homologs formed a distinct clade, but only contained homologs from the order Haloferacales (32.7% of the sequences).

The search for archaeal PPX homologs returned 96 sequences (**Figures 1C and S1**). Seven different orders were represented in the 92 sequences identified to at least this taxonomic level. The most represented were the Methanosarcinales, Sulfolobales, Methanobacteriales and Methanomicrobiales with 36.5%, 20.8%, 17.7% and 13.5% of the identified homologs respectively, while the remaining identified sequences were from the orders Methanomassiliicoccales (5.2%), Nitrososphaerales (1.0%) and Thermoplasmatales (1.0%). No PPX sequence was retrieved from the halophilic archaea.

Searches for other putative polyP synthesis (vacuolar transport chaperones 2 and 4^44^), or degradation (polyphosphate glucokinase and endopolyphosphatase^1,45^) proteins returned no significant hits in any archaeon.

### Physiological responses of M. mazei to Pi-starvation/overplus treatment

To further investigate anaerobic polyP accumulation in archaea, the methanogenic archaeon *M. mazei*, which harbours homologs to the main enzymes involved in polyP turnover (PPK1, PPK2 and PPX – **Figure 1**) was chosen for further study. *M. mazei* was grown in Pi replete medium for seven days to obtain sufficient cell biomass for the Pi-starvation/overplus experiment. Following inoculation in Pi-deficient media residual Pi in the medium was rapidly consumed by the cells to levels below the detection limit (2 µM) from day 2. During this period, the phosphate concentration in Pi-replete control was kept high (**Figure 2**). After 4 days, cells from Pi-starved cultures contained less than 2 µg P per mg protein. Upon the addition of Pi, there was an increase in the appearance of intracellular polyP inclusions within overplus cells (**Figure 3A**), compared both untreated (**Figure 3B**) and Pi-replete control (**Figure 3C**) cells. Overplus treated cells at day 6 contained an average of 0.223 µg of polyphosphate as Pi per mg of protein, compared to 0.066 µg polyphosphate as Pi per mg of protein in control samples (standard deviations ±0.090 and 0.145 respectively, p=0.0468). This represents a 237% increase in intracellular polyP levels in overplus cells previously exposed to Pi starvation.

**Figure 2.**
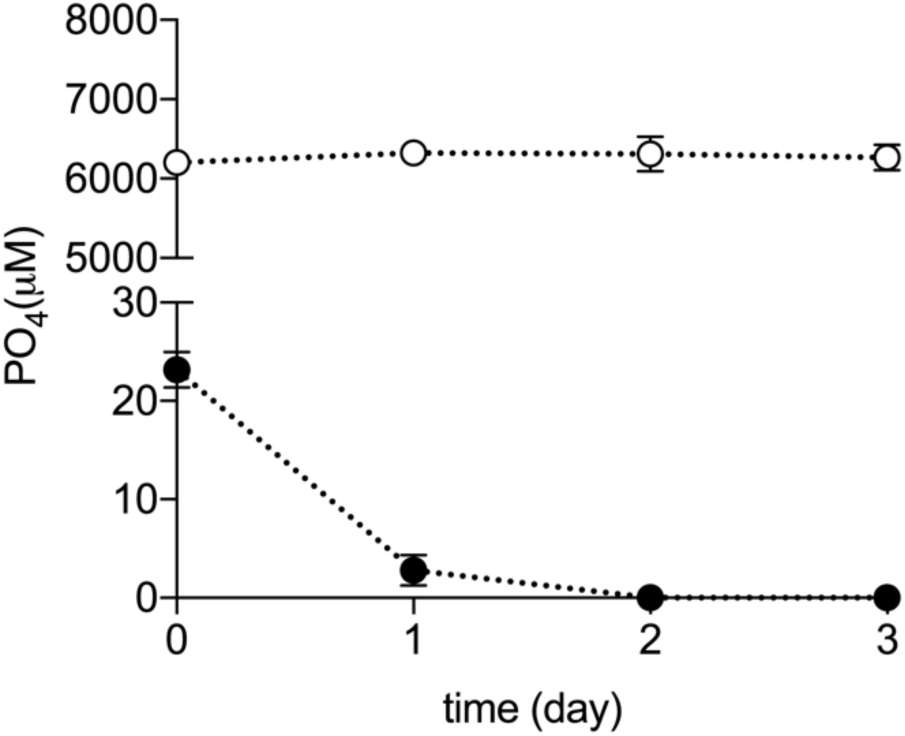
Change in extracellular phosphate concentration in *M. mazei* cultures under Pi-starved (filled circles) or Pi-replete (empty circles) conditions. Error bars indicate ±s.d. of biological triplicates (error bars may not be visible, due to small variability). Paired t-test indicated significant difference between treatment and control in all sampling days (p < 0.05).

**Figure 3.**
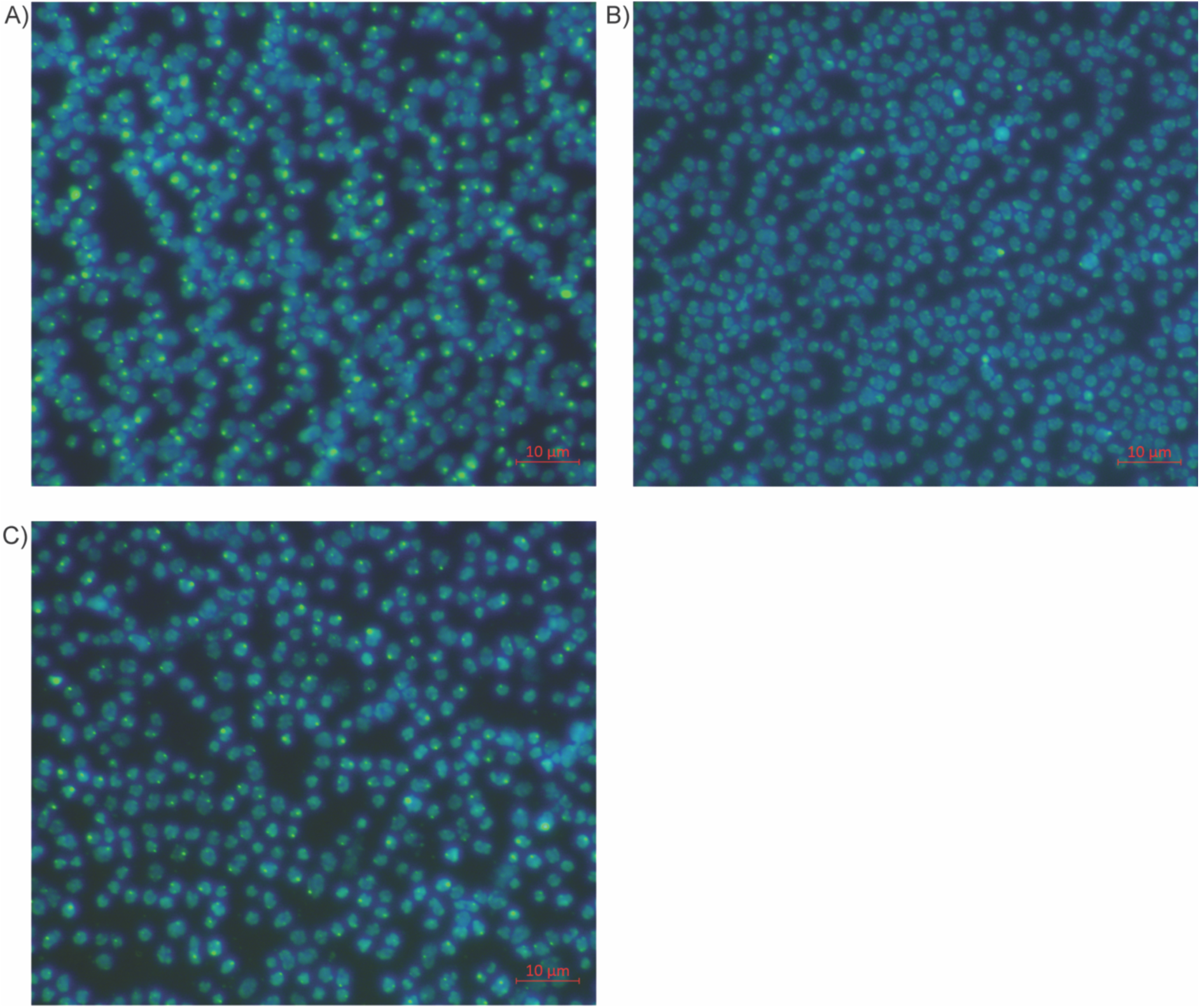
Epifluorescence microscopy of DAPI stained *M. mazei* cells: nucleic acids are visualised in blue, and polyphosphate granules in green. A) Cells sampled on day 6, following Pi-starvation/overplus treatment; B) *M. mazei* cells prior to Pi-starvation/overplus treatment; C) Cells from Pi-replete control on day 6. Images were captured with 1000X magnification. 10 µm reference bars are displayed on the bottom left corners.

Neither Pi depletion nor overplus significantly impacted methane production (**Figure 4A;** *α*=0.05). However, biomass production, as assessed by protein concentration, was significantly lower in Pi-starved cultures than in the controls (**Figure 4B**). This impact on the biomass protein was observed from day 2 onwards, coinciding with the beginning of the Pi depletion period (**Figure 2**).

**Figure 4.**
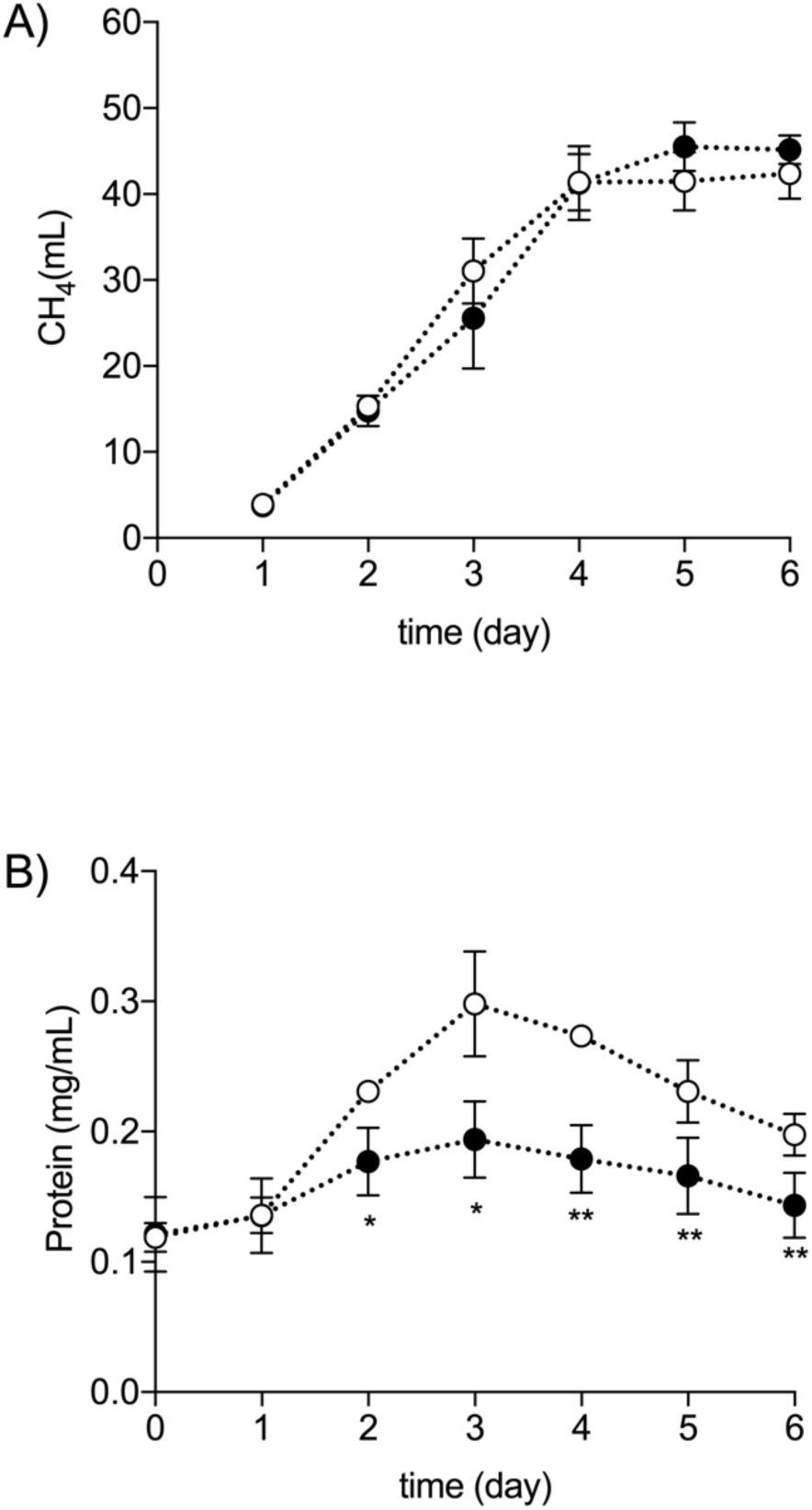
Growth, measured by the increase in protein concentration, and methane production by cells of *M. mazei* under conditions of either Pi-starvation or after overplus. A) Cumulative methane production. B) Protein concentration. Filled circles: Pi-starved/overplus; empty circles: Pi-replete control. Phosphate was added to the starved cultures on day 4. Error bars indicate ±s.d. of biological triplicates. Paired t-test was used to assess the difference between treatment and control in each sampling day. * p < 0.05; ** p < 0.01.

### Gene expression in response to changes in Pi availability

To determine the effect of Pi availability on the expression of *ppk1* and *ppk2*, we analysed transcript abundance in transcriptomic libraries prepared from cells of *M. mazei* both at the end of the Pi starvation phase (T4), and two days after Pi addition to the medium (the overplus phase, T6). In addition to *ppk1* and *ppk2*, we also evaluated changes in the expression of other genes known to be involved in cellular Pi turnover: *ppx*, the Pi starvation (PHO regulon) controlled genes *phoA* (alkaline phosphatase^46,47^), and those belonging to the high affinity ABC-type Pi transport complex (*pstSCAB-phoU*). The *pstSCAB-phoU* operon encodes the high affinity Pi binding protein PstS, the permeases subunits PstA and PstC, the cytoplasmic ATPase subunit PstB and the Pi transport rate modulator PhoU^48^.

Transcriptomes prepared from Pi starved *M. mazei* cells at day 4 were characterised by a large increase in the relative abundance of PHO regulated genes compared to the Pi replete control cells at the same time point. During this Pi starvation phase, the relative expression of *phoA* was 6.5-fold higher when compared to control Pi replete cultures (**Figures 5A and 6)**. Concomitant with the increased expression of *phoA*, we also observed 2.5-6.3-fold higher expression of the *pstSCAB-phoU* genes in Pi starved cells (**Figure 5A**). Genes from the two *pstSCAB-phoU* operons present in the genome of *M. mazei*^49^ were up-regulated. No significant difference (p<0.05) was observed in the relative abundance of either *ppk2* or *ppx* transcripts between starved and control cultures. The relative abundance of *ppk1* transcripts in Pi-starved cells was, however, 4.2-fold lower than in the Pi replete control cells (**Figures 5A and 6**).

**Figure 5.**
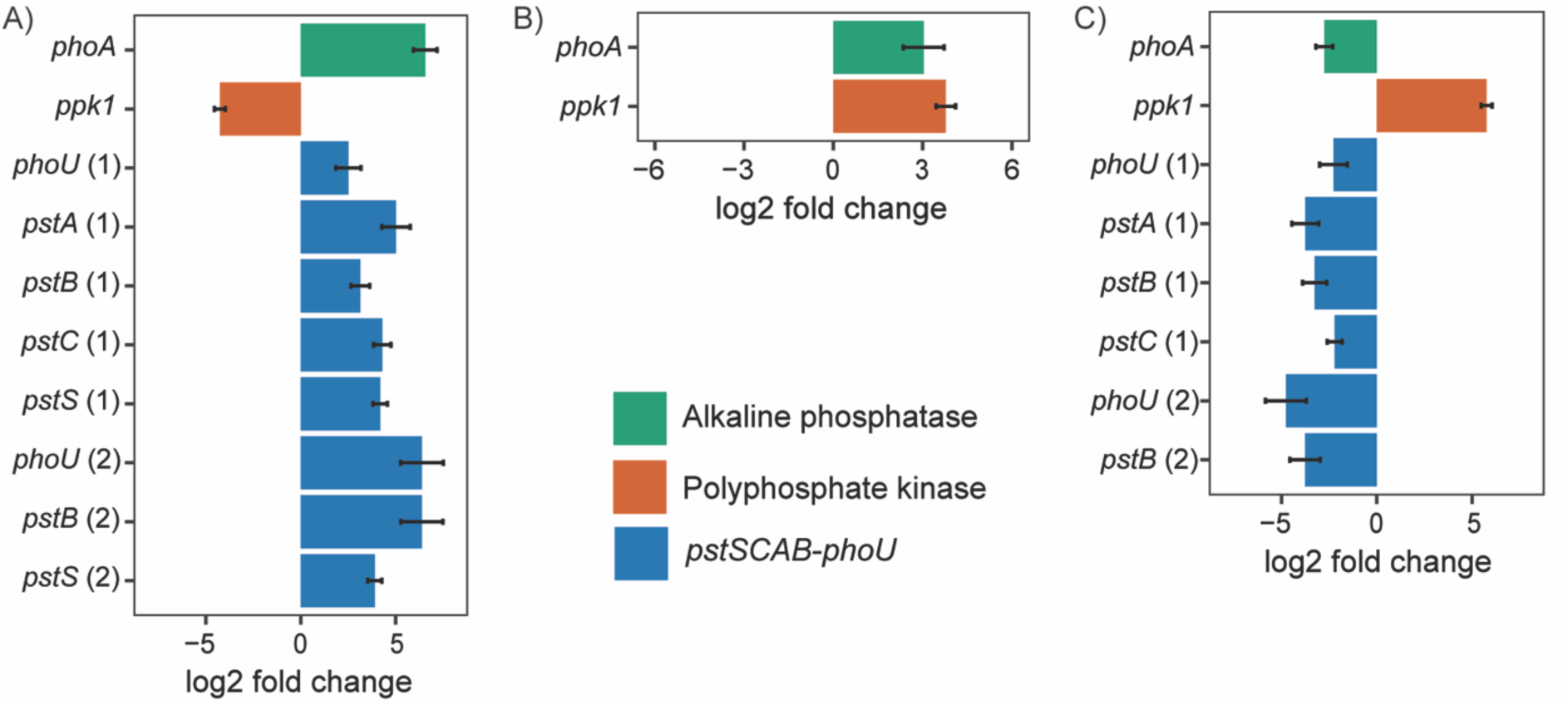
Differential expression analysis of *M. mazei* phosphate metabolism genes in response to Pi-starvation/overplus treatment. Bars < 0 indicate genes down-regulated, and > 0 indicate genes up-regulated in Factor 1 vs Factor 2: A) Pi-starved vs Pi-replete control at T4; B) Pi-overplus vs Pi-replete control at T6; C) Pi-overplus (T6) vs Pi-starved (T4). Only genes with fold change >2 and FDR< 0.05 were plotted. Numbers in brackets indicate the respective operon for the *PstSCAB-PhoU* complex.

**Figure 6.**
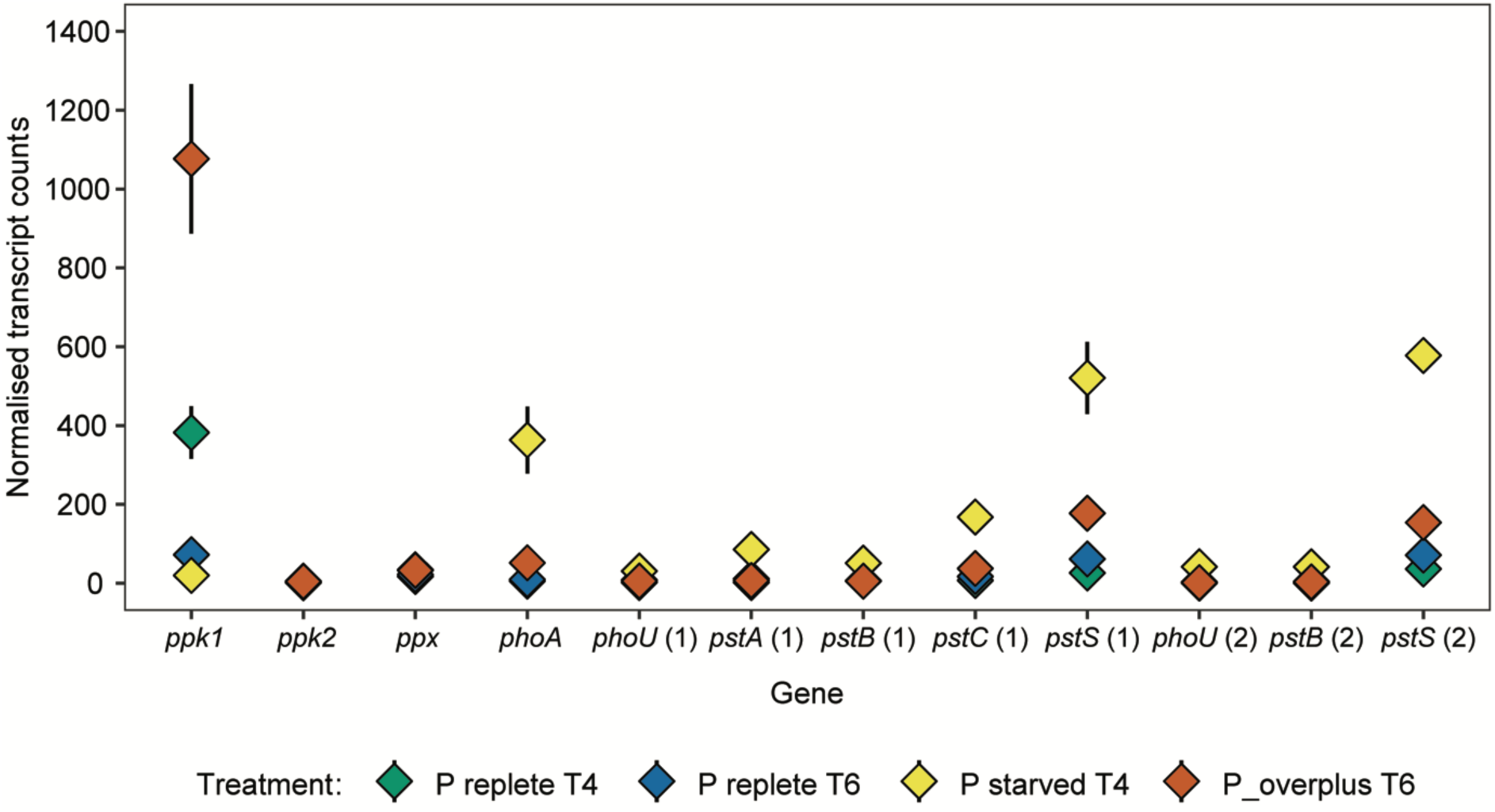
Normalised transcript counts from phosphate metabolism genes in *M. mazei* under Pi-starvation/overplus treatment. *ppk* 1/2, polyphosphate kinase; *ppx*, exopolyphosphatase; *phoA*, alkaline phosphatase; *phoU* and *pst* genes, components of the *PstSCAB-PhoU* complex. Numbers in brackets indicate the respective operon for the *PstSCAB-PhoU* complex.

The relief of Pi starvation during the overplus phase led to a large increase in the relative abundance of *ppk1* transcripts. When overplus cells of *M. mazei* at day 6 were compared to control cells not previously exposed to Pi starvation, at the same time point, the expression of *ppk1* was increased 3.7-fold (**Figures 5B and 6**). In comparison to Pi starved cells at day 4, the expression of *ppk1* in overplus cells was 5.7-fold higher (**Figures 5C and 6**). No significant difference (p<0.05) was observed in the relative abundance of either *ppk2* or *ppx* transcripts between the T6 overplus cells and either the T4 Pi starved or T6 Pi replete control cultures. Although the relative abundance of *phoA* was still significantly higher (3-fold) in overplus cells when compared to Pi replete control at day 6 (**Figure 5B**), the normalised transcript count was 50, in contrast to 363 detected during starvation at day 4 (**Figure 6**). Expression of the *pstSCAB-phoU* operon decreased significantly in overplus cells compared to cells under starvation: the relative abundance of *pstSCAB-phoU* transcripts was 2.5-4.7-fold less than that observed in Pi starved *M. mazei* cells (**Figure 5C**). There was no significant difference observed in *PstSCAB-phoU* expression between overplus cultures at day 6 and Pi replete control cells at the same time point.

In total, 474 genes exhibited a minimum two-fold expression change between Pi starved and control Pi replete cultures at T4 with 354 genes up- and 120 down-regulated. The imposition of overplus resulted in the differential expression of 446 genes, 419 of which were upregulated in comparison with the Pi-replete control and 27 downregulated at day 6.

With respect to in-treatment differences comparing overplus (day 6) with Pi-starved (day 4) cells, a total of 349 genes were classified as differentially expressed with 249 up- and 100 down-regulated. Gene ontology enrichment analysis assessed biological processes over-represented in both up- and down-regulated gene sets from cells under overplus conditions, in comparison to Pi-starved (**Figure S2**). In addition to polyP biosynthesis, several biological processes were represented among the up-regulated genes, including proteolysis and other catabolic pathways, response to stress, glycogen biosynthesis and nitrogen fixation. Regarding the down-regulated pathways, PHO regulated Pi metabolism was among the over-represented functions, along with different biosynthetic and transport processes.

## Discussion

Inorganic polyP metabolism is widely regarded as having a very ancient evolutionary origin^50^, and this is supported by the presence of genes related to its metabolism in organisms from all three domains of life^13,51,52^. However, the prevalence and function of polyP in archaea is poorly understood. Searching the NCBI RefSeq database for the known enzymes involved in polyP turnover found 442 putative PPK1, PPK2 and PPX homologs from diverse taxa across the Archaea Domain, although it remains to be determined whether these homologs have activity. Whilst putative PPK1 homologs were detected in diverse members of the class Halobacteria and PPK2 homologs in the order Haloferacales, no PPX homologs were found across the class Halobacteria. By contrast, while PPX homologs were identified within the Crenarchaeota, concentrated in the order Sulfolobales, no PPK1 or 2 homologs were retrieved from this taxon. Martinez-Bussenius *et al*.^53^ has previously shown the presence of *ppx* but not *ppk1* in 6 species of *Sulfolobus* sp., and *Metallosphaera sedula*. Wang *et al*.^22^ using proteomic analysis have also shown the lack PPK1 or 2 in *M. sedula* and *Sulfolobus sp.* Our analysis showing no PPK1 or PPK2 like sequences across all 5 Orders of Crenarchaeota within RefSeq would suggest that the lack of polyP-synthesising genes is a feature of the Crenarchaeota phylum. Furthermore, homologues to PPX are present only in the Crenarchaeota Order Sulfolobales: the known enzymes for polyP turnover would thus appear to be absent from the other 4 Orders of Crenarchaeota. It is possible that as yet uncharacterised pathways are used by members of the taxa Halobacteria and Sulfolobales (and the Crenarchaeota) for polyP degradation and synthesis, respectively. Investigation of fully sequenced and annotated genomes, and of representative cultures from within these taxa is, however, required to address this question. Previous studies have shown continued polyP accumulation in bacterial *ppk* mutants leading to speculation that other enzyme systems exist for polyP synthesis^54,55^. PPK1, 2 and PPX homologs were found throughout the orders Methanomicrobiales, Methanosarcinales and Methanomassiliicoccales, suggesting that these methanogenic archaea may cycle polyP in a manner more similar to traditional bacterial metabolism than in the Halobacteria or Crenarchaeota.

The addition of Pi to cells previously subjected to Pi starvation induces the rapid accumulation of polyP in a phenomenon known as “polyP overplus” (reviewed in ref.^56^). Although the accumulation of polyP in response to extreme oscillations in Pi availability has been demonstrated in both bacteria and yeasts, such Pi starvation induced polyP formation is poorly explored in archaea and has not hitherto been demonstrated under anaerobic conditions. In this paper, we describe intracellular accumulation of polyP by the methanogenic archaeon *M. mazei* upon exposure to overplus conditions. This organism displayed the full known genotype for polyP accumulation, which represented the profile observed for the majority of the methanogens retrieved from the database. The increase in intracellular polyP accumulation by overplus-treated cells (**Figure 3**) was accompanied by a 5.7-fold increase in *ppk1* transcript abundance by day 6 when compared to the same Pi-starved cultures on day 4. No increase in *ppk2* or *ppx* expression was observed in the control or overplus cells at both sample points. When compared to the control cells at the end of the starvation phase (T4) the expression of *ppk1* was 4.2-fold lower in the treated cells, rising to 3.7-fold higher at after overplus (T6). Upregulation of *ppk1* expression would therefore appear to begin after the addition of Pi to Pi-starved *M. mazei* cells and thus differ from the majority of previous bacterial overplus studies where *ppk* (and *ppx*) expression has been shown to begin during the Pi starvation phase, under the control of the PHO regulon^57–63^. Nevertheless, a Pi starvation phase is required to upregulate ppk1, as no increase in *ppk1* expression was observed in Pi replete control cells at day 6. The expression of *ppk1* is, therefore apparently not induced by Pi starvation in *M. mazei* but is primed by it.

The PHO regulon describes those Pi starvation-inducible genetic elements involved in P assimilation. Under conditions of Pi limitation, the upregulation of PHO controlled genes, such as the *pstSCAB-phoU* operon, and the alkaline phosphatase gene *phoA* occurs via a two-component sensor-regulator complex consisting of the proteins PhoB and PhoR^57^. *M. mazei* cells, under conditions of Pi limitation do elicit a PHO activation response with an increase in the expression of both the *pstSCAB-phoU* operon, and *phoA* under conditions of Pi limitation. Although the PHO regulon is distributed across all kingdoms of life, its activation by Pi limitation within Archaea has not been widely studied, being previously limited only to the halophilic aerobic archaeon *Halobacterium salinarum*^64^. In *M. mazei*, nitrogen limitation has also been shown to cause up-regulation of *pstS* and *phoU*^65^, which was attributed to a possible high demand of Pi under nitrogen starvation. In another study with *M. mazei*, high salinity induced the expression of genes for the *pstSCAB-phoU* complex^8^. Although polyP was not assessed, the authors suggested that the up-regulation of the Pi transport system could be accompanied by polyP production, and possibly associated with a regulatory role of this molecule under conditions of high salinity. *M. mazei* is found in environments often experiencing Pi scarcity, such as freshwater and marine sediments^66^. The strong response of these genes under Pi starvation suggests that they are closely regulated by environmental changes and that *M. mazei* might be adapted to respond quickly to Pi oscillations, which may confer increased fitness.

The mechanisms involved in *ppk1, ppk2* and *ppx* regulation within cells of *M. mazei*, and in the Archaea more generally, are at present unknown. Non PHO regulon based models for the transcription of these polyP metabolism genes have, however, been proposed in bacteria and yeasts whereby a complex network of interactions involving stress induced proteins such as RelA, SpoT, RpoS, ppGpp, and cellular sigma factors coordinate transcription^1,63^. Further studies are thus required to determine if similar regulatory networks exist within *M. mazei* with respect to the regulation of *ppk1, ppk2* and *ppx*. Hereof, it is noteworthy that a previous study has suggested the absence of ppGpp in Crenarchaeota^67^, highlighting that regulatory differences may occur between the Domains Bacteria and Archaea. The triggers that induce expression of *ppx* (and thus allow *M. mazei* cells access to intracellular polyP reserves), and the physiological function of PPK2, also require further investigation.

Despite the impact on cellular biomass, Pi starvation did not affect methane production significantly. In fact, the energetics underpinning polyP accumulation remains puzzling, and it is intriguing that organisms living on the edge of the thermodynamic limit^68^ invest in a high energy demand process^16^. In addition, other biosynthetic processes known to require large quantities of energy, such as nitrogen fixation and glycogen synthesis, were also up-regulated simultaneously with polyP production. Elucidating the polyP energetics and functions in methanogens may reveal mechanisms related to the responses of these organisms to Pi availability, contributing to the understanding of the eco-physiology of these organisms with a paramount role in global carbon cycling.

Although polyP metabolism under anaerobic conditions has, for the most part, been associated with its degradation, for example within the engineered Enhanced Biological P Removal process (where polyP utilization takes place during anaerobiosis with other biopolymer formation occurring upon subsequent exposure to aerobic or anoxic conditions^69–71^), or naturally in some sediments, polyP accumulation under anaerobic conditions is not without precedent. For example, Szulmajster and Gardner^72^ observed intracellular polyP accumulation in a *Clostridia* sp. during dissimilative growth on creatinine, whilst Rudnick *et al.*^16^ demonstrated the presence polyP in cells of *M. mazei* (previously named *M. frisia*) when grown on either methanol or H_2_/CO_2_, and Liang *et al.*^73^ have shown that *Rhodopseudomonas palustris* accumulates polyP under anaerobic illuminated conditions. PolyP granules were also visualised in the hyperthermophilic sulphate-reducing archaeon *Archaeoglobus fulgidus*^74^. Additionally, Keating *et al.*^75^ reported the presence of polyP granules in the microbial biofilm present in an anaerobic digester performing P removal during wastewater treatment. These results, coupled with our own observations of *M. mazei*, provide further evidence that not only can polyP formation occur under anaerobic conditions, but that it can do so in response to nutritional stress. Furthermore, our sequence analysis would suggest that the potential for polyP metabolism is much more widespread amongst the Archaea than previously thought. Given that aerobic bacterial polyP metabolism has been shown to play an important role in both the marine^76^ and freshwater^77^ P cycles - particularly under varying external Pi concentrations - the contribution of both archaeal and anaerobic microbial communities to global P cycling processes requires further consideration. Additionally, understanding the triggers which promote both polyP accumulation - and thus Pi removal - and its degradation could lead to the development of optimized Pi recovery systems for application within anaerobic digester units.

## Acknowledgments

We thank Nicola Byrne for initial support with the cultures, Simon Isaksson for technical support, and Dr. Alyona Minina for assisting F.S.P with microscopy.

The Financial support of Science Foundation Ireland and the Northern Ireland Department of Education and Learning, through grant 14/IA/2371, is gratefully acknowledged.

## Author Contributions

J.M., V.O.F., F.A., K.A.M., P.M. and J.P.Q. wrote the grant proposal; F.S.P, J.Ch., A.S., J.M. and V.O.F. designed the experiments; J.Ch. performed the phylogenetic analysis and polyP quantification; F.S.P, A.S. and J.Co. performed culture experiments; F.S.P and B.M. carried out transcriptomic analysis and the data was analysed by F.S.P and N.W; F.S.P, J.Ch. and J.M wrote the manuscript. All authors critically reviewed the manuscript.

## Competing Interests

The authors declare no competing interests.

## Additional Information

Sequencing data has been deposited in the Sequence Read Archive of the National Center for Biotechnology (accession numbers SAMN08132863 - SAMN08132874).

Supplementary information accompanies this paper.

